# Differential detection of *Tomato mosaic virus* (ToMV) and *Tomato brown rugose fruit virus* (ToBRFV) using CRISPR-Cas12

**DOI:** 10.1101/2021.03.16.435580

**Authors:** Dan Alon, Hagit Hak, Menachem Bornstein, Gur Pines, Ziv Spiegelman

## Abstract

CRISPR/Cas12-based detection is a novel approach for efficient, sequence-specific identification of viruses. Here we adopt the use of CRISPR/Cas12a to identify the *Tomato brown rugose fruit virus* (ToBRFV), a new and emerging *Tobamovirus* causing substantial damage to the global tomato industry. Specific guide RNAs (gRNAs) were designed to detect either ToBRFV or the closely related *Tomato mosaic virus* (ToMV). This technology enabled the differential detection of ToBRFV and ToMV. Sensitivity assays revealed that viruses can be detected from 15-30 ng of RT-PCR product, and that specific detection could be achieved from a mix of ToMV and ToBRFV. In addition, we show that this method enabled the identification of ToBRFV in samples collected from commercial greenhouses. These results demonstrate a new method for species-specific detection of plant viruses. This could provide a platform for the development of efficient and user-friendly ways to distinguish between closely related strains and resistance-breaking pathogens.

## Main text

Plant viruses of the *Tobamovirus* genus (family: Virgaviridae) are important crop pathogens, causing significant damage to the global agriculture industry (Dombrovsky et al. 2017; Oladokun et al. 2019; Broadbent 1976). The *Tobamovirus* genus contains several Solanaceae-infecting species, including *Tobacco mosaic virus* (TMV) and *Tomato mosaic virus* (ToMV). Tobamovirus particles are rod-shaped, encapsulating a single-stranded sense RNA (+ssRNA) genome of approximately 6.4kb encoding four ORFs. ORFs 1 and 2 encode the two subunits of the viral replicase complex, separated by a read-through stop codon. ORF3 encodes the viral 30 kDa movement protein (MP). ORF4 encodes the 17-18 kDa viral coat protein (CP) (Dawson et al.1986; Goelet et al. 1982). Tobamoviruses are highly infectious and transmitted by mechanical contact with working hands, tools, soil, and other infected plants (Reingold et al. 2016; Tomlinson 1987; Panno et al. 2020).

*Tomato brown rugose fruit virus* (ToBRFV), a new tomato-infecting tobamovirus emerged in tomato greenhouses in Israel and Jordan in 2014 (Salem et al. 2016; Luria et al. 2017). ToBRFV was found to overcome all tobamovirus resistance genes in tomatoes, including the durable *Tm-2^2^* resistance gene, which has remained unbroken for over 60 years (Luria et al. 2017). Outbreaks of ToBRFV in Europe (Beris et al. 2020; Alfaro-Fernández et al. 2020; van de Vossenberg et al.2020; Skelton et al. 2019; S. Panno et al. 2019; Menzel et al. 2019), North America (Camacho-Beltrán et al. 2019; Ling et al. 2019), and Asia (Yan et al. 2019; Fidan et al. 2019) indicate a rapidly emerging global epidemic.

Serological methods such as enzyme-linked immunosorbent assay (ELISA) and western-blot are currently used to detect ToBRFV (Luria et al. 2017; Klap et al. 2020a; Klap et al. 2020b;Levitzky et al. 2019). However, these methods lack species-specificity and cannot be used to distinguish between ToBRFV and closely related tobamoviruses such as TMV and ToMV. This is due to the high conservation of the tobamovirus CP, resulting in antibody cross-reactivity between tobamovirus species (Luria et al. 2017). To cope with this challenge, protocols for specific detection of ToBRFV were developed including deep sequencing (Luria et al. 2017), sequence-specific reverse-transcription polymerase chain reaction (RT-PCR) primers (Luria et al. 2017), Real-time RT-PCR (Stefano Panno et al. 2019) and loop-mediated isothermal amplification (LAMP) (Sarkes et al. 2020).

An emerging approach for the precise identification of viruses is based on CRISPR/Cas technology. The CRISPR/Cas is an innate immune system present in many bacteria and archaea (Jiang and Doudna 2017). In this system, Cas proteins target specific viral sequences by the formation of a ribonucleoprotein (RNP) complex with a single guide RNA (sgRNA) complementary to the invading DNA sequence (Gasiunas et al. 2012). Cas proteins then cleave the bound invading nucleic acid sequence, thereby neutralizing the pathogen (Charpentier and Marraffini 2014). While this system is mostly utilized for genome editing (Jinek et al. 2012; Hwang et al.2013; Shalem et al. 2014; Garst et al. 2017), more recently it has been shown that some Cas proteins can be used for the detection of specific nucleic acid sequences (Chen et al. 2018;Harrington et al. 2018; Gootenberg et al. 2017). For example, Cas12 obtains a non-specific single-stranded DNAse activity following specific interaction with its DNA substrate (Chen et al. 2018). This property was used by Chen and colleagues to apply Cas12 as a biosensor: in addition to the RNP complex of *Lachnospiraceae bacterium* Cas12a and its gRNA, a fluorophore quencher (FQ)–labeled single-stranded DNA substrate was added. Upon Cas12 activation, it non-specifically degrades ssDNA, releasing the fluorophore from its quencher, resulting in a fluorescent signal. This approach was performed for the detection of human papillomavirus and has also proven efficient in SARS-Cov-2 detection (Chen et al. 2018; Broughton et al. 2020) demonstrating the great potential of this method in viral diagnostics.

Recently, CRISPR/Cas12 was applied for the detection of TMV, *Potato virus Y* (PVY), and *Potato virus X* (PVX) (Aman et al. 2020), suggesting its potential application in the detection of plant viruses. Here we adopt the CRISPR/Cas12 technology to specifically identify the emerging ToBRFV and to distinguish it from a closely related tobamovirus, ToMV.

To determine the ability of CRISPR/Cas12a to distinguish between ToBRFV and ToMV, we have set up an experimental design based on the conserved features and the variation between these two viruses (Fig. 1). A region within the tobamovirus ORF1 was used for the design of species-specific gRNAs (Fig. 1, A). The CRISPOR platform (Concordet and Haeussler 2018) was used to design four ToMV-specific gRNAs (*tomv-1, tomv-7, tomv-8*, and *tomv-9*) and three ToBRFV-specific gRNAs (*tobrfv-3, tobrfv-5*, and *tobrfv-9*), based on the ToMV and ToBRFV genomes (GenBank accession nos. AF332868.1 and KX619418.1, respectively). The resulting gRNAs were concatenated to a T7 promoter sequence (Supplementary table 1), and gRNAs were transcribed using TranscriptAid T7 High Yield Transcription Kit (Thermo Scientific) and purified using Zymo RNA Clean & Concentrate kit (Zymo).

**Fig. 1.**
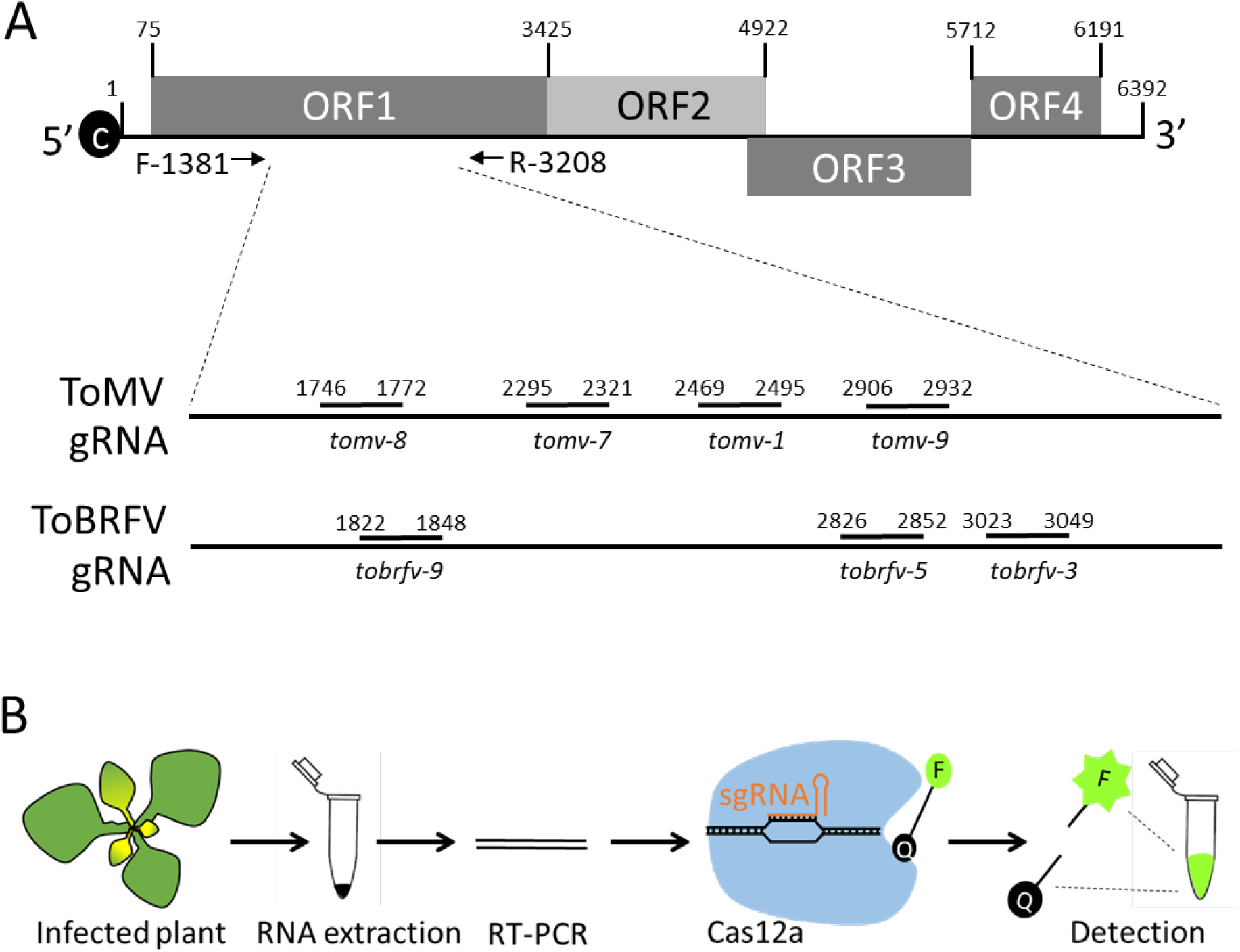
Differential detection of ToMV and ToBRFV using CRISPR/Cas12a. **A,** A 1827 bp fragment was targeted for differential detection. Four gRNAs were designed for ToMV detection (*tomv-1, tomv-2, tomv-8* and *tomv-9*) and three gRNAs were designed for ToBRFV detection (*tobrfv-3, tobrfv-5* and *tobrfv-9*). B, Illustration of plant virus detection using CRISPR/Cas12. Leaf samples are collected from the infected plant. RNA is extracted from the infected leaf and RT-PCR is performed. Using species-specific gRNA, Cas12a-gRNAs specifically targets the sequence of ToBRFV or ToMV. Activation of Cas12a by sequence-specific binding triggers the degradation of the ssDNA probe and releases the fluorophore (F) from the quencher (Q) to emit the fluorescent signal.

To test the ability of the different gRNAs to detect ToMV or ToBRFV, 3-week old tomato plants (*Solanum lycopersicum* L. cv. Moneymaker) (LA2706) were inoculated separately with ToMV and ToBRFV. After three weeks, RNA was extracted from newly emerged leaves showing viral symptoms using the Plant Total RNA Mini Kit (Geneaid, RPD050), and cDNA was synthesized using the qPCRBIO cDNA Synthesis Kit (PCRBIO, PB30.11). The ORF1 region was then amplified using the Q5^®^ High-Fidelity 2X Master Mix (NEB) with primers F (5’-ccaggtctgagtgggatg-3’), and R (5’-gtctcaccttgtacctcatgtac-3’) (Fig. 1, B). PCR products were purified using Zymo DNA Clean & Concentrator kit (Zymo).

For CRISPR/Cas12 detection, gRNA-LbCas12a (NEB) complexes were prepared by mixing 62.5nM gRNA with 50nM LbCas12a in NEBuffer 2.1 (NEB) to a final volume of 20uL and incubated at 37°C for 30 minutes (Fig. 1, B). Next, 1uM ssDNA FAM reporter (/56-FAM/TTATTATT/3BHQ_1/) and 1nM (120 ng) of RT-PCR products were added to the complexes and incubated for 10 minutes at 37°C. Cleavage of the ssDNA FAM reporter and release of the fluorophore from the quencher was measured using Tecan Spark plate-reader with an excitation wavelength of 485nm, and emission was measured at 535nm (Fig. 1, B).

We tested the different sgRNA to specifically identify each virus in the CRISPR/Cas12 fluorescence assay. For ToMV detection, all four *tomv* sgRNA-Cas12a complexes were incubated with the ToMV RT-PCR product. The ToBRFV RT-PCR product served as a negative control (Fig. 2, A). Among these sgRNAs, *tomv-1* and *tomv-2* emitted a robust fluorescent signal in response to ToMV, which was 4.2- and 2.5-times higher than with the ToBRFV negative control (Fig. 2, A and B). The *tomv-8* sgRNA produced a fluorescent signal 57% higher than the control, and *tomv-9* signal was similar in ToMV and ToBRFV (Fig. 2, A). A reciprocal experiment was performed for ToBRFV detection, only this time with the ToMV RT-PCR product served as a negative control (Fig. 2, D). Here, *tobrfv-3* produced a strong fluorescent signal when incubated with the ToBRFV template, which was 6.5 times higher than the ToMV control template (Fig. 2, C, and D). The *tobrfv-5* and *tobrfv-9* sgRNAs were also efficient in ToBRFV detection, producing a fluorescent signal that was 5- and 1.9-higher than the ToMV control (Fig. 2, C). These results established the ability of the CRISPR/Cas12a for species-specific identification of ToMV and ToBRFV. Since the most efficient detection of ToMV and ToBRFV was done by the *tomv-1* and *tobrfv-3 s*gRNAs, respectively, these sgRNAs were selected for further analysis.

**Fig. 2.**
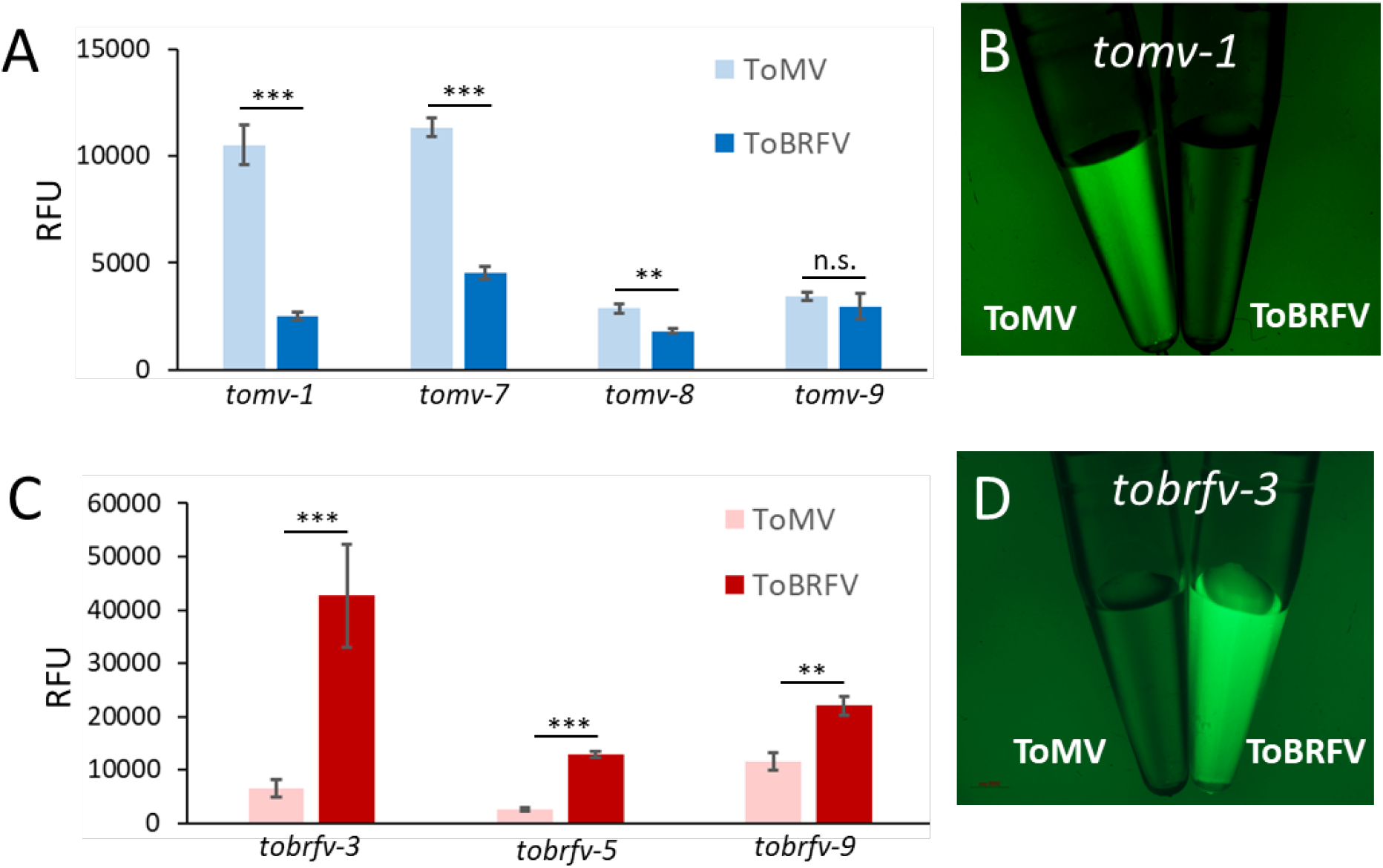
Analysis of different sgRNAs for the detection of ToMV and ToBRFV. A, CRISPR/Cas12a-based fluorescent detection of ToMV using the different gRNAs on samples from ToMV- (light blue) and ToBRFV- (dark blue) infected tomato plants. B, Fluorescence image of ToMV detection using the *tomv-1* gRNA. C, CRISPR/Cas12a-based fluorescent detection of ToBRFV using the different gRNAs on samples from ToMV- (pink) and ToBRFV- (red) infected tomato plants. D, Fluorescence image of ToBRFV detection using the *tobrfv-3* gRNA. RFU=relative fluorescence units. n.s= not significant. **P≤0.01, ***P ≤0.001 in Student’s t-test.

To test the sensitivity of this system, a series of dilutions was performed for each of the ToMV and ToBRFV RT-PCR products (Fig. 3 A and B). While 15 ng Rt-PCR products were sufficient for detecting ToMV (Fig. 3, A), 30ng of RT-PCR product were needed for the detection of ToBRFV (Fig. 3, B). Since 15-30 ng of PCR products were the minimal threshold for detection, we further used these parameters to examine if the CRISPR/Cas12a system can specifically detect each virus in the case of a mixture of ToMV and ToBRFV. To test this, detection assays were performed on a mixture of RT-PCR products from both ToMV and ToBRFV. Notably, 15 ng of ToMV were detected in a mixture with 15 ng of ToBRFV, resulting in a fluorescent signal that was 85% higher than the signal obtained with 30 ng of ToBRFV only (Fig. 3, C). In addition, the CRISPR/Cas12a system was also able to detect 15 ng of ToBRFV mixed with 15 ng of ToMV, producing a fluorescent signal that was 3.5-fold higher than the signal received with only 30 ng of ToMV (Fig. 3, D). These results suggest that the CRISPR/Cas12a system can be used to detect ToBRFV, even in the case of mixed infection with the related virus ToMV.

**Fig. 3.**
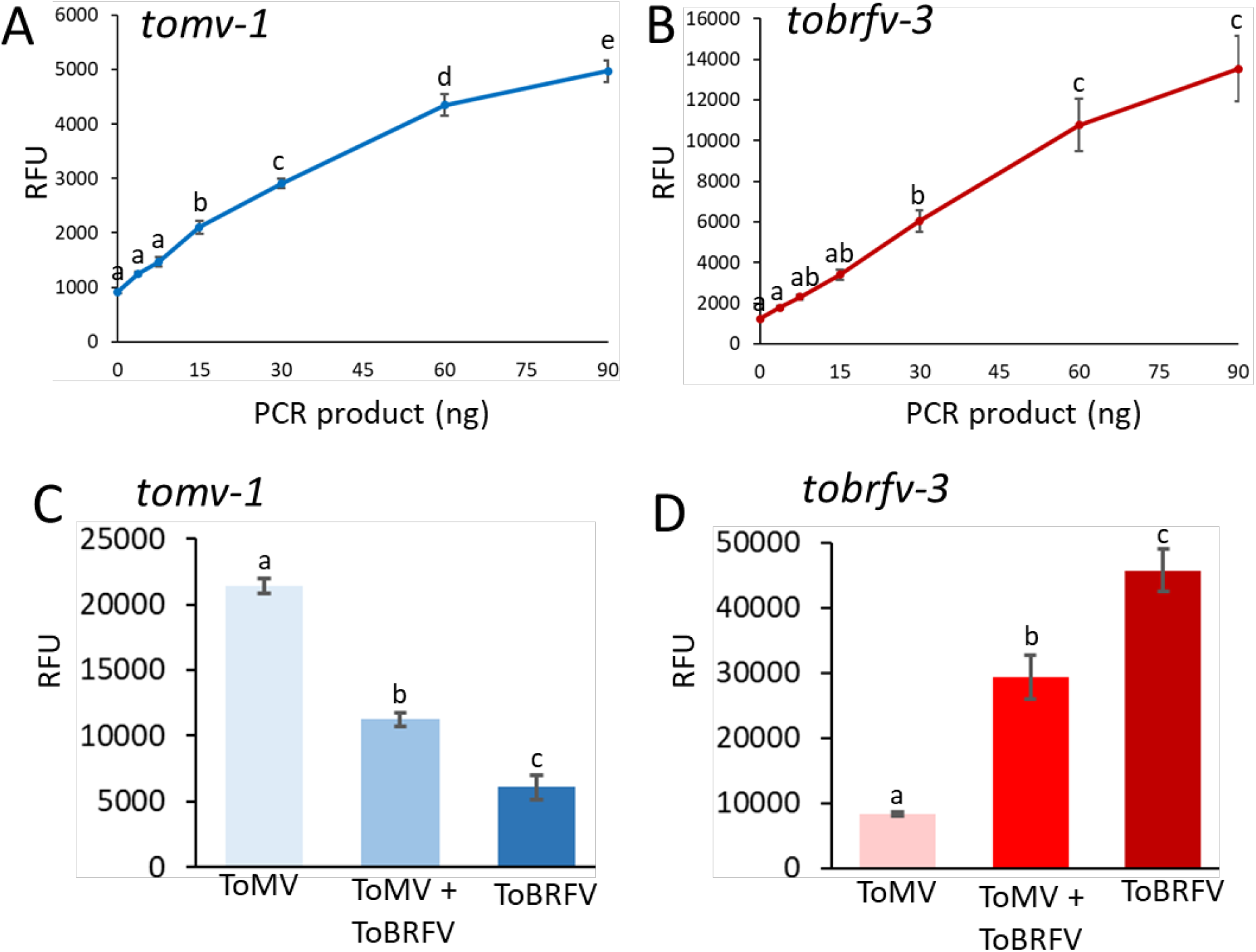
Sensitivity and specificity analysis for the selected sgRNAs. A, Detection of ToMV using the *tomv-1* gRNA in a series of dilutions of ToMV RT-PCR products. B, Detection of ToBRFV using the *tobrfv-3* gRNA in a series of dilutions of ToBRFV RT-PCR products. C, Identification of ToMV in a mix of ToMV and ToBRFV RT-PCR products from using the *tomv-1* gRNA. D, Identification of ToBRFV in a mix of ToMV and ToBRFV RT-PCR products using the *tobrfv-3* sgRNA. Different letters indicate statistical significance in Tukey-HSD test (P≤0.05).

Next, we tested the applicability of the CRISPR/Cas12a system for specific detection of tobamoviruses in field samples. In December 2020, tomato plants with mosaic symptoms were detected in greenhouses of two growers in Azriel village, Israel (Fig. 4, A). We used our CRISPR/Cas12a system to diagnose if the plants were infected with ToMV or ToBRFV. Analysis using ToMV gRNA *tomv-1* revealed that both tomato plants are likely not infected with ToMV, as indicated by the weak fluorescent signal that was similar to the ToBRFV control (Fig. 4, B). In marked contrast, the ToBRFV gRNA *tobrfv-3* showed high fluorescence for both samples, which were similar to the ToBRFV positive control (Fig. 4, C). To confirm these results, RT-PCR products from both samples were sequenced. Indeed, both samples showed 100% identity to ToBRFV-IL (KX619418.1) (Fig. 4, D). These results establish that the CRISPR/Cas12 system can detect ToBRFV in field samples.

**Fig. 4.**
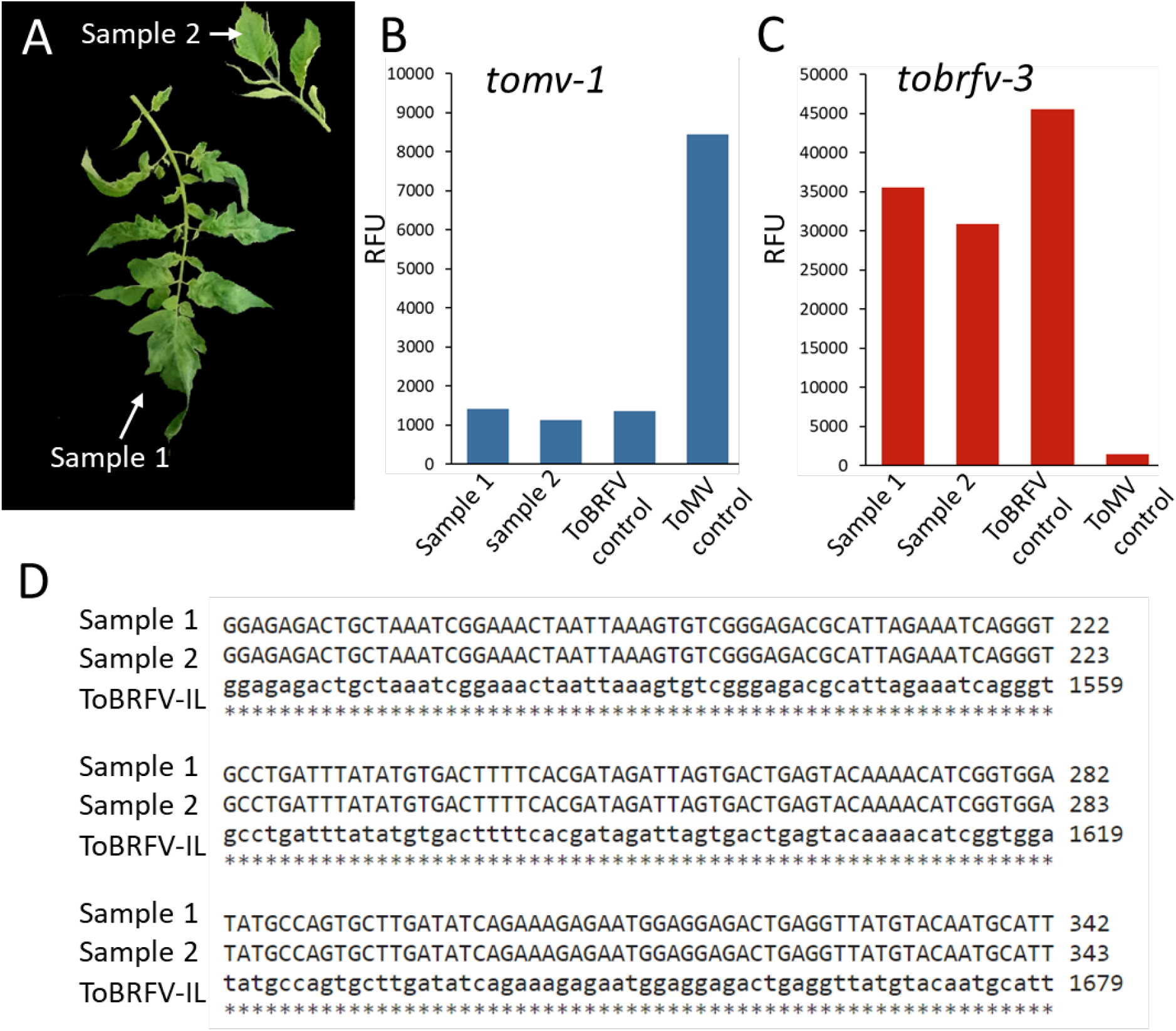
Detection of ToBRFV in field samples using CRISPR/Cas12a. A, Tomato with mosaic leaf pattern were detected in two different sites in Azriel village, Israel, in December 2020. Samples were obtained from two infected tomato cultivars, Ikram and Sgula. B, CRISPR/Cas12a-based detection using the *tomv-1* gRNA indicated no presence of ToMV. C, CRISPR/Cas12a-based detection using the *tobrfv-3* gRNA indicated the presence of ToBRFV in the sample. D, Sequencing results from the two samples established the presence of ToBRFV in the sample.

ToBRFV is an emerging tobamovirus, which is spreading rapidly throughout the world and poses a substantial threat to the global tomato industry. ToBRFV symptoms are similar to other tobamoviruses such as ToMV, and conventional serological detection methods are currently unable to fully differentiate between ToBRFV and other members of the tobamovirus genus. These challenges drove the development of new Real-time PCR and LAMP assays for specific ToBRFV detection (Stefano Panno et al. 2019); (Sarkes et al. 2020)). While it was shown that CRISPR/Cas12a can be used to detect plant viruses (Aman et al. 2020), it was unknown if it can be applied to distinguish between closely related plant viruses of the same genus. Here, we provide evidence that the CRISPR/Cas12a technology can also be applied to identify ToBRFV and distinguish it from the closely related ToMV (Fig. 2). Using this method, ToBRFV can be detected with low concentrations of RT-PCR products, and also when mixed with ToMV (Fig. 3). Detection of ToBRFV in field samples (Fig. 4) demonstrates this technology’s potential as a *bonafide* virus identification method that can be used in agricultural practice.

Further research can enhance the CRISPR/Cas12a method for more accessible and user-friendly applications. For example, CRISPR/Cas12a can be combined with isothermal amplification methods such as loop-mediated isothermal amplification (LAMP) or recombinase polymerase amplification (RPA), to detect ToBRFV quickly, without the need for a thermocycler, establishing an accurate technology that would not require complex hardware or methodology. Recently, it was shown that the amplification step may be avoided through signal amplification. One such option is to use gold nanoparticles, or L-Methionine gold nanoclusters combined with electrochemiluminescence to enhance and stabilize the signal (Choi et al. 2020; Liu et al. 2021). Another approach was recently reported where the use of several gRNAs, targeting several loci within the target DNA, significantly increase the fluorescent signal achieved following Cas13 activation (Fozouni et al. 2020). In addition, several reports have shown that the output signal can be visual rather than fluorescent, further simplifying the detection procedure. Such outputs may be colorimetric, or by using lateral flow strips, similar to common pregnancy tests (Choi et al.2020; Broughton et al. 2020). Collectively, these approaches will allow onsite, rapid, and accurate detection without the need for specialized training or equipment. These techniques will be especially relevant for the detection of closely related pathogen species or strains, and resistance-breaking variants.

## Acknowledgements

We would like to thank Ms. Neta Mor (Extension Service, Israeli Ministry of Agriculture and Rural Development), for providing tomato field samples and Dr. Victor Gaba (ARO, Israel) for his critical reading of this manuscript. This work was funded by grant number 20-02-0130 from the Chief Scientist, Israeli Ministry of Agriculture.

## Author contributions

DA, HH, GP and ZS designed the research. DA, HH and MB conducted the experiments. GP and ZS wrote the manuscript.

---

**Supplementary Table 1.**
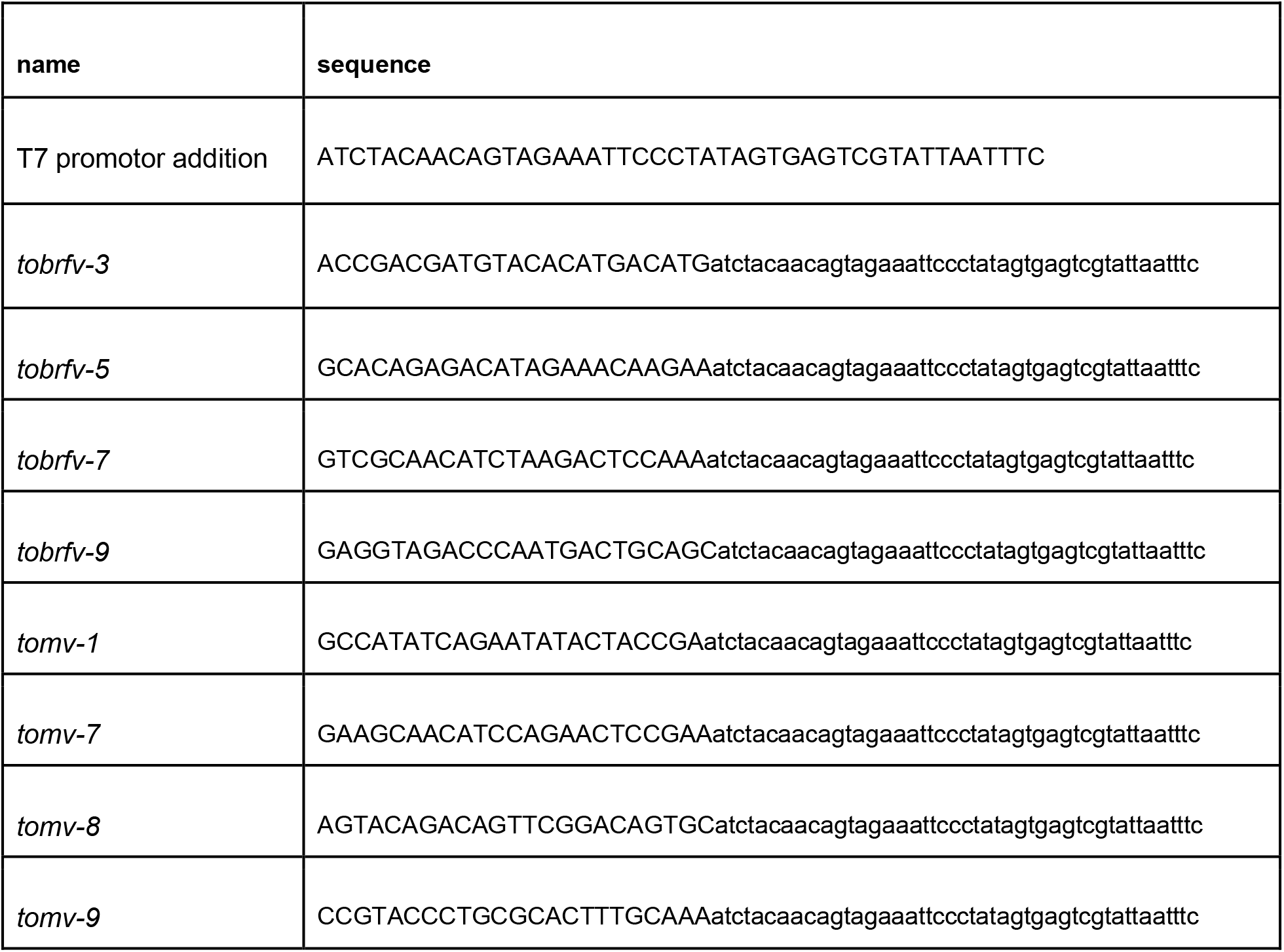
Oligonucleotides used in this manuscript

